# Bayesian tip dating reveals heterogeneous morphological clocks in Mesozoic birds

**DOI:** 10.1101/350496

**Authors:** Chi Zhang, Min Wang

**Affiliations:** Key Laboratory of Vertebrate Evolution and Human Origins, Institute of Vertebrate Paleontology and Paleoanthropology, Chinese Academy of Sciences, Beijing 100044, China; Center for Excellence in Life and Paleoenvironment, Chinese Academy of Sciences, Beijing 100044, China

**Keywords:** Mesozoic birds, tip dating, relaxed clock, MrBayes

## Abstract

Recently, comprehensive morphological datasets including nearly all the well-recognized Mesozoic birds become available, making it feasible for statistically rigorous methods to unveil finer evolutionary patterns during early avian evolution. However, few quantitative and statistical studies have yet been performed. Here, we exploited the advantage of Bayesian tip dating under relaxed morphological clocks to infer both the divergence times and evolutionary rates while accounting for their uncertainties. We further subdivided the characters into six body regions (i.e., skull, axial skeleton, pectoral girdle and sternum, forelimb, pelvic girdle, and hindlimb) to assess evolutionary rate heterogeneity both along the lineages and across partitions. We observed extremely high rates of morphological character changes during early avian evolution and the clock rates are quite heterogeneous among the six regions. The branch subtending Pygostylia shows extremely high rate in the axial skeleton, while the branches subtending Ornithothoraces and Enantiornithes show very high rates in the pectoral girdle and sternum, and moderately high rates in the forelimb. The extensive modifications in these body regions largely correspond to refinement of the flight capability. The rest of the relatively slow and even rates suggest that there is no dominant selective pressure in favoring of modifications in the skull and pelvis. This study reveals the power and flexibility of Bayesian tip dating implemented in MrBayes to investigate evolutionary dynamics in deep time.

## Introduction

Birds are one of the most speciose (over 10,000 recognized species) and ecological diverse living vertebrates (Gill 2007). The origin and evolution of birds have long been a hot debate in evolutionary biology, although it has been generally accepted that birds underwent two large-scale radiations during their 160 million years evolution, one for the stem groups in the Cretaceous, and the other for the crown groups in the Paleogene (Mayr 2009; Prum et al. 2015; Wang and Zhou 2017). Over the last few years, numerous well-preserved Mesozoic bird fossils have been described (Chiappe and Meng 2016; Wang and Zhou 2017), and consensus has been approached regarding their systematic relationships (O’Connor and Zhou 2013; Wang et al. 2017; 2018). These wealthy data have significantly bridged the large morphological gap between birds and their non-avian theropod predecessors (O’Connor and Zhou 2015; Wang and Zhou 2017), and demonstrated their evolutionary success related to novel traits. Expanded morphological characters that cover the major Mesozoic avian groups with chronological data become accessibly recently (Wang and Lloyd 2016), making it possible to trace the early avian evolution more quantitively and to address important questions such as the divergence times of the major clades and the patterns of morphological character changes in and between lineages (Brusatte et al. 2014; Lloyd 2016). However, few quantitative and statistical studies for early avian evolution have yet been performed.

In a previous study, Wang and Lloyd (2016) investigated evolutionary rate heterogeneity in Mesozoic birds under maximum parsimony using a large morphological dataset containing 262 characters and 58 taxa (Wang et al. 2015). The approach was stepwise: first to infer the most parsimonious trees, then to inform the internal node ages using certain *ad hoc* measures, and last to obtain the branch rates from dividing the number of parsimonious changes by the time durations along the corresponding branches. It did not account for the uncertainties of tree topology, times and rates statistically, and was unable to model the evolutionary process explicitly. The dataset was then extended to 280 characters and 68 taxa recently (Wang and Zhou 2018), with newly recognized species and significantly revised morphological character scorings. Moreover, we revised this dataset according to (Field et al. 2018), mainly focusing on the cranial morphology of Ichthyornithiformes and Hesperornithiformes. The modified dataset contains nearly all the well-recognized Mesozoic birds and represents the most comprehensive morphological characters at present, thus provides more power to unveil finer evolutionary patterns and becomes applicable to more statistically rigorous methods.

We further subdivided the characters into six partitions, each representing a different anatomical region (i.e., skull, axial skeleton, pectoral girdle and sternum, forelimb, pelvic girdle, and hindlimb), to assess evolutionary rate heterogeneity. Partitioned analysis has been attempted by Clarke and Middleton (2008) on a much smaller dataset of stem birds under a Bayesian non-clock model to examine the branch-length variations across three body regions. Since each branch length was a product of geological time duration and evolutionary rate, the two elements could not be distinguished without a time tree and clock assumption.

Here, we exploited the advantage of tip dating to infer both the divergence times and evolutionary rates while accounting for their uncertainties in a coherent Bayesian statistical framework. The technique was originally developed for analyses combining both morphological and molecular data (Pyron 2011; Ronquist et al. 2012a; 2016; Zhang et al. 2016; Lee 2016; Gavryushkina et al. 2016) and has been termed as “total-evidence dating”. It has also been productively applied to morphological data only (Lee et al. 2014; Bapst et al. 2016; Matzke and Wright 2016; King et al. 2017) and we use “tip dating” for that matter. This approach has the essential strengths of incorporating various sources of information from the fossil record directly in the analysis, modeling the speciation process explicitly through a probabilistic model allowing for parameter inference and model selection, and utilizing the state-of-the-art developments in Bayesian computation.

## Materials and methods

### Morphological data

The morphological data used in this study is based on Wang and Zhou (2018) which is extended and revised from Wang et al. (2015). Character scorings for *Ichthyornis dispar, Hesperornis regalis, Parahesperornis alexi*, and *Baptornis advenus* were further revised according to Field et al. (2018). The modified dataset consists of 280 morphological characters and 68 operational taxonomic units (Dromaeosauridae as the outgroup, 65 Mesozoic and 2 extant birds). Detailed character descriptions are listed in Supplementary Material.

The characters were further partitioned into six anatomical regions to assess evolutionary rate heterogeneity across these regions: skull (53 characters), axial skeleton (36 characters), pectoral girdle and sternum (48 characters), forelimb (65 characters), pelvic girdle (23 characters), and hindlimb (55 characters). Each partition included more than 20 characters to ensure sufficient information for inference.

### Tip dating

In Bayesian tip dating framework, we infer the posterior probability distribution of the model parameters, which combines the information from the morphological characters (likelihood) and the priors (including the distributions of the fossil ages and the other parameters in the tree model and clock model). For the likelihood, the Mkv model (Lewis 2001) was used for the character state substitution with gamma rate variation across characters. The gamma distribution has mean 1.0 (*α* = *β*) and was approximated with four discrete categories (Yang 1994). 36 characters were defined as ordered (Supplementary Information), which means instantaneous change is only allowed between adjacent states, the rest of the characters were thus unordered. The prior for gamma shape *α* was exponential(1), while the priors for the time tree and the relaxed clock model parameters are described in detail in the following sections.

In order to root the tree properly and infer the evolutionary rates more reliably, we applied five topology constraints as Aves, Pygostylia, Ornithothoraces, Enantiornithes and Ornithuromorpha (Fig. 1), each of which forms a monophyletic clade.

**Figure 1.**
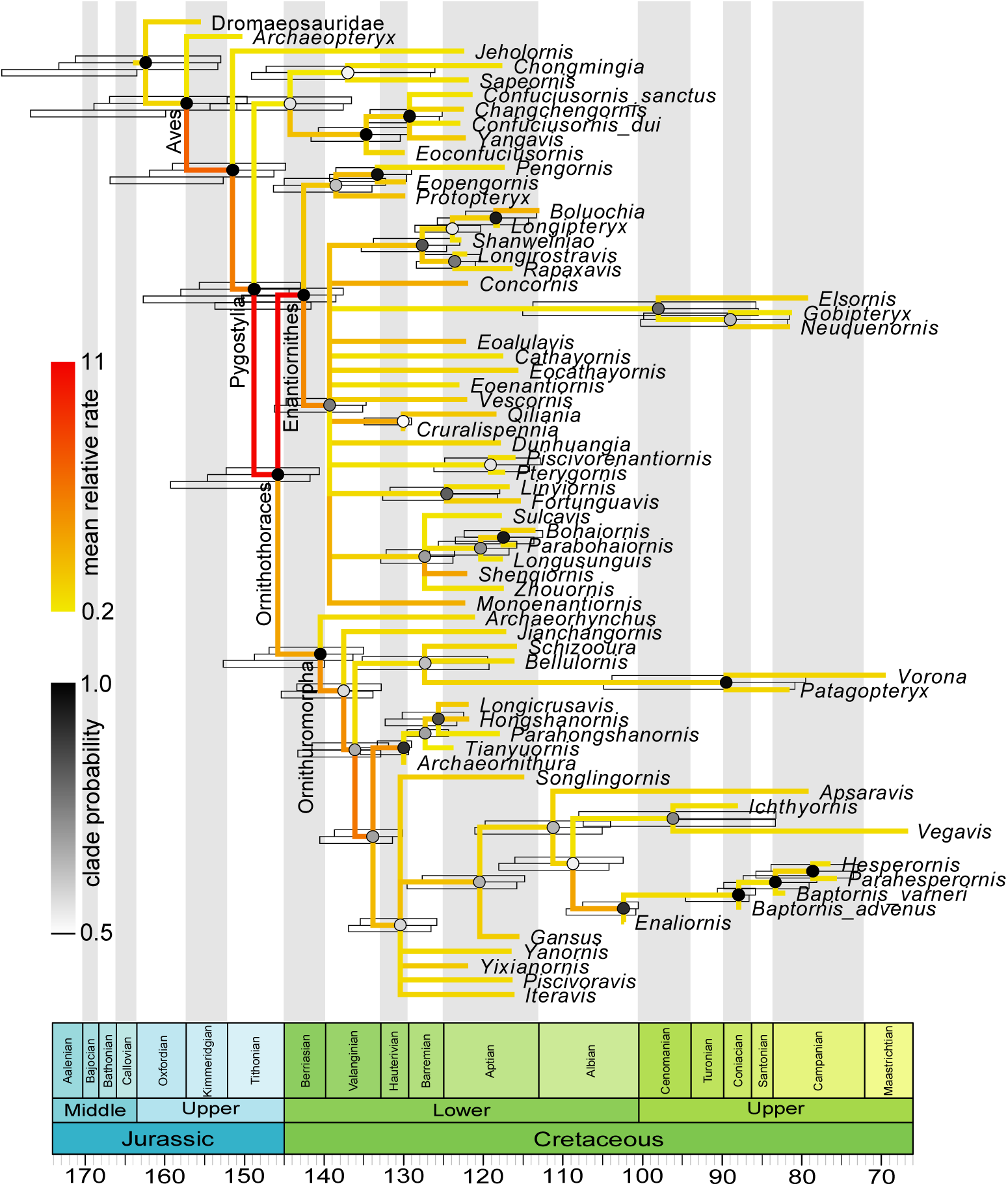
Dated phylogeny (time tree) of Mesozoic birds. The topology (majority-rule consensus tree) shown is inferred allowing fossil ancestors (*r* = 0) without partitioning the morphological characters. The node ages in the tree are the posterior medians and the shade of each circle represents the posterior probability of the corresponding clade. The color of the branch represents the mean relative clock rate at that branch. The error bars on the top (in blue) at the internal nodes denote the 95% HPD intervals of age estimates. In comparison, the error bars below (if present, in cyan) denote the 95% HPD intervals when disallowing fossil ancestors (*r* = 1) under a single partition. Additionally, the error bars shown at the early avian diversifications (in green) are the 95% HPD intervals of age estimates when the characters are partitioned into six anatomical regions and disallowing fossil ancestors (*r* = 1). The two extant species (*Anas* and *Gallus*) were included in the analyses but not shown in the representation (as a sister clade of *Vegavis*).

The posterior distribution was estimated using Markov chain Monte Carlo (MCMC). We executed two independent runs with four chains (one cold and three hot) per run for 40 million iterations and sampled every 2000 iterations. The first 25% samples were discarded as burn-in for each run, and the remaining samples from the two runs were combined after checking consistency between runs.

### Tree model

The fossilized birth-death process (Stadler 2010; Heath et al. 2014; Gavryushkina et al. 2014; Zhang et al. 2016) was used to model speciation, extinction, fossilization and sampling, which gave rise to the prior distribution of the time tree 𝒯. The process starts at the root with two lineages sharing the same origin. Each lineage bifurcates with a constant rate *λ* and goes extinct with a constant rate *μ*. Concurrently, each lineage is sampled with a constant rate *ψ* and is removed from the process upon sampling with probability *r*. Extant taxa are sampled with probability *ρ*. The explicit derivation of the probability density function was given in Gavryushkina et al. (2014, Equation 4). When the removal probability 0 ≤ *r* < 1, the sampled tree may contain fossil ancestors (i.e., fossils with sampled descendants), while setting *r* = 1 disables fossil ancestors (i.e., all fossils are at the tips).

The age of each fossil bird was assigned a uniform prior with lower and upper bounds from the corresponding stratigraphic range (Supplementary Information). The root age was assigned an offset exponential prior with mean 169 Ma (slightly older than the first appearance datum of Dromaeosauridae) and minimum 153 Ma (slightly older than the first appearance datum of *Archaeopteryx*). For inference, we reparametrized the speciation, extinction and sampling rates and assigned priors as *d* = *λ* – *μ* – *rψ* ∼ exponential(100) with mean 0.01, *v* = (*μ* + *rψ*)/*λ* and *s* = *ψ*/(*μ* + *ψ*) ∼ uniform(0, 1). The sampling proportion of extant taxa (*Anas* and *Gallus*) was set to 0.0002, based on the number of described living bird species around ten thousands (Gill and Wright 2006).

### Clock model

We applied the independent gamma (white noise) relaxed clock model (Lepage et al. 2007) to investigate evolutionary rate heterogeneity both along the tree and across the six anatomical regions (partitions). The model was reparametrized aiming to articulate the relative rates. Specifically, the model assumes that the substitution rate (clock rate, in unit of substitutions per character per myr) of branch *i*, in partition *j c*_*ij*_, is a product of the mean rate *c* and the relative rate *r*_*ij*_ (i.e., *c*_*ij*_ = *cr*_*ij*_), and *r*_*ij*_ is gamma distributed with mean 1.0 and variance *σ*_*j*_/(*t*_*i*_*c*), where *t*_*i*_ is the geological time duration of branch *i* (*i* = 1, …, 2*m* – 2, *j* = 1, …, *n*). Thus, the clock model has *n* + 1 parameters (*c, σ*_1_, …, *σ*_*n*_), and there are (2*m* − 2) *× n* independent rates in the tree. The relative rate *r*_*ij*_ serves as a multiplier to the mean rate. Large deviation of *r*_*ij*_ from 1.0 indicates severe heterogeneity of the morphological clock, while *r*_*ij*_’s all being similar to 1.0 models a somewhat strict clock. The branch length (distance, in unit of substitutions per character) in the Mkv likelihood calculation, *b*_*ij*_, is the product of time duration *t*_*i*_ and clock rate *c*_*ij*_ (i.e., *b*_*ij*_ = *t*_*i*_*c*_*ij*_ = *t*_*i*_*cr*_*ij*_).

The prior used for the mean clock rate *c* was gamma(2, 200) with mean 0.01 and standard deviation 0.007, and that for *σ*_*j*_ was exponential(10).

## Results and Discussion

There is no clear evidence for us to believe that all fossils are at the tips *a priori*, however, we encountered severe mixing problem in the MCMC when allowing fossil ancestors while partitioning the morphological characters into six anatomical regions. When treating the characters as a single partition on the other hand, we were able to achieve good mixing both allowing and disallowing fossil ancestors (setting *r* = 0 and *r* = 1 respectively). Thus, we first show the results without partitioning the data and compare the difference between with and without fossil ancestors, then we focus on the evolutionary rate heterogeneity when the data is partitioned (only under *r* = 1). In general, the parameter estimates are quite similar whether fossil ancestors are allowed, the difference is more dramatic whether the data is partitioned (see below).

### Single partition

The phylogeny estimated from tip dating allowing fossil ancestors (*r* = 0) is shown in Figure 1. The tree is well resolved, with a few polytomies mainly nested within Enantiornithes that represent the uncertainty of the taxa relationships. This topology agrees with previously published trees in the placements of the major clades (Wang et al. 2015; Wang and Lloyd 2016; Wang and Zhou 2018). The root age is estimated at 162.56 (153.00, 171.26) Ma (see also Table 1), which covers the fixed age of 168.7 Ma (1 myr older than the first appearance datum of Dromaeosauridae using the minimum branch length method) (Wang and Lloyd 2016). The posterior age of Dromaeosauridae is 154.90 (141.66, 167.69) Ma, mainly within Late Jurassic, while the prior range expands the whole Cretaceous (66.0, 167.7). As the posterior mean relative rate at the branch of Dromaeosauridae is 1.08 (close to 1.0), the similarity of the morphological characters informs a short time span. The mean ages of the divergences of Pygostylia, Ornithothoraces and Enantiornithes are about 6–8 myr older than the fixed ages in the previous study (Wang and Lloyd 2016). We emphasize that the age estimates using tip dating integrates all available sources of information but the minimum branch length method only used the first appearance datum of the oldest taxa thus might underestimate the ages.

**Table 1.** Posterior distributions (mean and 95% HPD interval) of model parameters

| | single partition, $r = 0$ | single partition, $r = 1$ | six partitions, $r = 1$ |
| --- | --- | --- | --- |
| $\alpha$ | 1.81 (1.35, 2.31) | 1.82 (1.35, 2.31) | 1.80 (1.33, 2.27) |
| $d = \lambda - \mu - r\psi$ | 0.010 (0.00087, 0.022) | 0.010 (0.00075, 0.021) | 0.010 (0.00042, 0.020) |
| $\nu = (\mu + r\psi)/\lambda$ | 0.98 (0.94, 1.00) | 0.93 (0.81, 1.00) | 0.92 (0.79, 1.00) |
| $s = \psi/(\mu + \psi)$ | 0.028 (0.00047, 0.075) | 0.33 (0.00051, 0.90) | 0.35 (0.00059, 0.90) |
| $t_{\text{mrca}}$ | 162.56 (153.00, 171.26) | 164.17 (153.33, 173.34) | 172.61 (164.01, 180.81) |
| $c$ | 0.012 (0.0090, 0.015) | 0.011 (0.0083, 0.014) | 0.010 (0.0081, 0.012) |
| $\sigma$ or $\sigma_1$ | 0.058 (0.035, 0.084) | 0.058 (0.036, 0.084) | 0.070 (0.026, 0.12) |
| $\sigma_2$ | | | 0.069 (0.028, 0.12) |
| $\sigma_3$ | | | 0.057 (0.028, 0.090) |
| $\sigma_4$ | | | 0.053 (0.025, 0.087) |
| $\sigma_5$ | | | 0.042 (0.0017, 0.084) |
| $\sigma_6$ | | | 0.061 (0.029, 0.095) |

When disallowing fossil ancestors (*r* = 1), one concern might arise is that some node ages might be overestimated due to forcing every tip to be the result of speciation. However, the difference from allowing fossil ancestors (*r* = 0) is really minor in our case. The node ages are only about 1–2 myr older if not otherwise similar (Fig. 1). The topologies in these two cases are almost identical, except for two places—one is the placement of *Cruralispennia* which becomes unresolved in the big polytomy within Enantiornithes, the other is *Archaeornithura* which becomes a sibling of *Tianyuornis*. In fact, the estimated proportion of fossil ancestors is 0.13 (0.06, 0.21), indicating most fossils are indeed tip fossils. *Enaliornis* and *Archaeornithura* have the highest posterior probabilities of being ancestral (0.95 and 0.93 respectively).

The evolutionary rates are thus very similar for the two cases and we only show the result under *r* = 0 (Fig. 1). The mean clock rate (*c*) is estimated around 0.01 substitutions per character per myr (i.e., approximately one character-state change per 100 million years) (Table 1). The relative clock rate at each branch represents the deviation from the mean rate. We observe extremely high rates during early avian evolution (Fig. 1). The relative rates at the two branches from Aves to Pygostylia are 6.63 (0.97, 17.49) and 5.34 (0.01, 15.52), and accelerate to 10.89 (0.00, 32.5) at the branch subtending Ornithothoraces. High rates are also encountered along the early branches of Ornithuromorpha, then slow down substantially towards crownward branches including the one leading to extant birds. Enantiornithes shows even higher rate of 11.24 (1.49, 30.75) when it diverges from Ornithuromorpha in the Early Cretaceous, and the rates decrease dramatically in its later history. These observations concurs with previous comparative studies that birds underwent a large scale of diversification in tandem with the dinosaur-bird transition (Benson and Choiniere 2013; Lee et al. 2014; Wang and Lloyd 2016).

### Six partitions

The evolutionary rates inferred above are averaged across all morphological characters. Further partitioning the data into six anatomical regions make it feasible for us to estimate refined evolutionary rates both along branches and across partitions. Different partitions have their own independent rate variations while sharing the same tree topology and geological time duration (i.e., single time tree 𝒯). As mentioned above, the mixing was very poor if fossil ancestors were allowed (*r* = 0) and independent runs did not give consistent estimates. The difficulty was probably due to limited data in each partition interfered with inefficient reversible-jump MCMC (rjMCMC) algorithm (Green 1995) implemented. Further investigations are required. At the moment, we just show the result disallowing fossil ancestors (*r* = 1) which simplifies the tree structure with no need for rjMCMC. In this case, the major clades inferred in the tree are unchanged, although a few taxa with large uncertainties shuffle a bit (Supplementary Figs S1–S6). Comparing with the node ages estimated under a single partition, those at the early avian diversifications are slightly older (Fig. 1, see also Table 1) while the younger ages become more similar. The age differences are more dramatic whether the data is partitioned than whether fossil ancestors are allowed.

The rates of morphological character changes are quite heterogeneous among the six regions during early avian evolution (Fig. 2), although the mean rate (*c*) estimated is almost identical as before (Table 1). The branch subtending Pygostylia shows extremely high rate in the axial skeleton (Fig. 2, Supplementary Fig. S2), which is one order of magnitude higher than in the rest five regions. Clearly, the high rate observed here indicates extensive morphological changes in the vertebral column, and the most recognizable change is that a long tail consisting of over 20 caudal vertebrae in *Archaeopteryx* and *Jeholornis* is replaced by a short element called pygostyle which is formed by the fusion of several caudal most vertebrae (Wang and Zhou 2017). The transition from long to short tail is functionally important for the evolution of powered flight in birds: a short tail could forward the gravitational center, and with attached feathers become indispensable for the avian flight apparatus (Gatesy and Dial 1996). The branches subtending Ornithothoraces and Enantiornithes show very rapid morphological changes in the pectoral girdle and sternum, and moderately high rates in the forelimb (Fig. 2, Supplementary Figs S3&S4). In comparison, the corresponding rates in the other regions are close to 1.0 with slight variation. These results suggest that most of the changes related to the shoulder and forelimb are close to the origin of Ornithothoraces, for example, the presence of an ossified sternum with a keel (attachment for the major flight muscle in modern birds) and further elongate forelimb (O’Connor and Zhou 2015), contributing to the refinement of flight capability. The rapid morphological changes towards Enantiornithes correspond to their unique shoulder morphology relative to the ancestor Ornithothoraces; for instance, enantiornithines have a sternum with a caudally restricted keel and an elongate acromion of the scapula, both of which are the major components of the flight apparatus in birds (Chiappe and Walker 2002; Wang and Zhou 2017). Previous morphometric study suggested that Enantiornithes have different flight style compared with other Mesozoic birds in terms of limb proportion (Dyke and Nudds 2009). The rest of the relatively slow and even rates suggest that there is no dominant selective pressure in favoring of modifications in the skull and pelvis.

**Figure 2.**
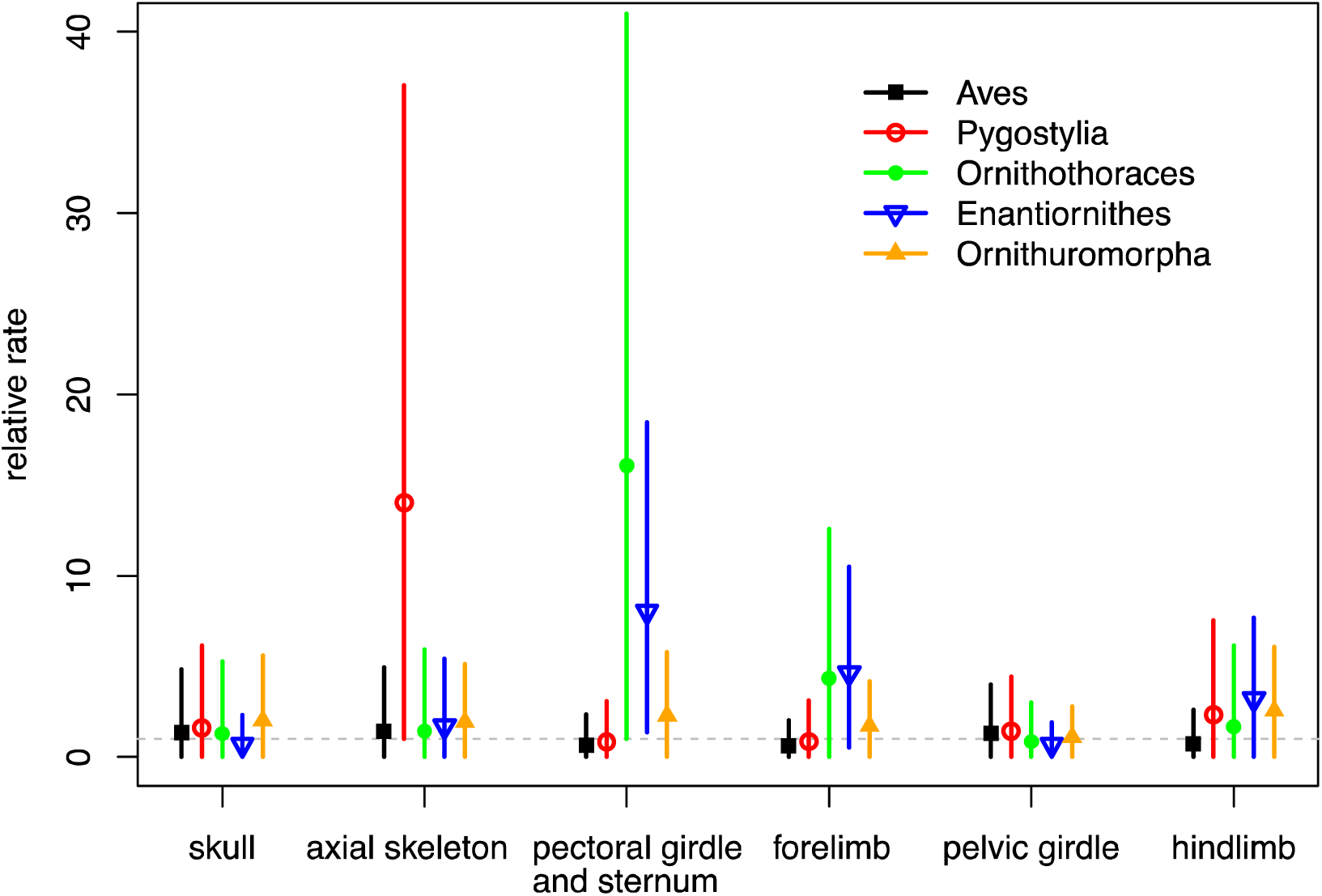
Posterior estimates of the relative clock rates along the branches subtending the major transitions of early avian evolution for six anatomical regions of the bird body. The dot and error bar denote the mean and 95% HPD interval for each estimate. The horizontal dashed line indicates the mean relative rate of 1.0 in the relaxed model.

When we compared the evolutionary rates between Enantiornithes and Ornithuromorpha across the six regions by summarizing the mean relative rates within each clade (excluding *Anas, Gallus* and their common ancestral branch), the differences are not striking, with medians uniformly close to 1.0 (Fig. 3). However, significantly high rates (outliers) are detected along some early diverging branches within these two clades, and a slowdown in more crownward branches (Fig. 3, Supplementary Figs S1–S6), suggesting that morphological changes are rapid in early diversifications and the process slows down subsequently due to saturated ecological niches for these two avian groups, as proposed in previous study (Wang and Lloyd 2016).

**Figure 3.**
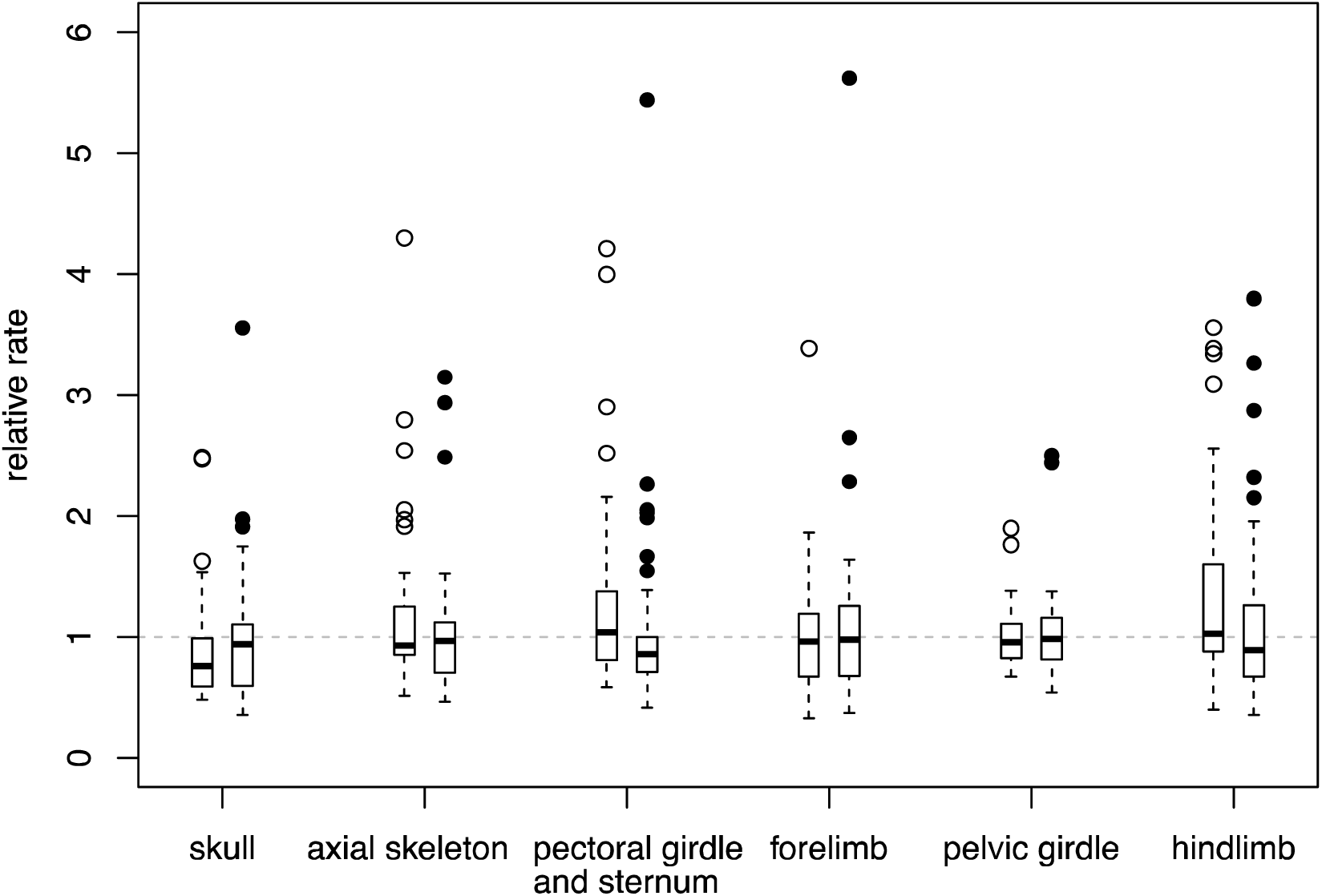
Boxplots summarizing the mean relative clock rates across branches in the Enantiornithes clade (left) and Ornithuromorpha clade (right) respectively, for the six anatomical regions. The box denotes the 1^st^, 2^nd^ (median) and 3^rd^ quartiles while the dots are the outliers. The horizontal dashed line indicates the mean relative rate of 1.0 in the relaxed clock model.

To test the robustness of age estimates to the root age prior and the impact on evolutionary rates in consequence, we halved and doubled the range between the mean and minimal in the original prior, that is, using offset-exp(153, 161) and offset-exp(153, 185) priors for the root age. This comparison showed slightly varied posterior age estimates deep in the tree. For the root age in particular, the estimates are 170.83 (161.33, 179.78) and 174.31 (164.34, 184.77) under the smaller and larger prior mean respectively. Thus, the conclusion of evolutionary rate heterogeneity above is not changed as the age differences are minor comparing with the significantly high rates which are usually one order of magnitude higher than 1.0.

Another concern related to the accuracy of age estimates might be the nonuniform fossil record both in its geographical and stratigraphic distribution. Although more complicated models allowing fossil sampling rate vary over time are available (Gavryushkina et al. 2014; Zhang et al. 2016), it is not practical to apply in our case to make reliable inference. The current model assuming constant sampling rate is a balance between model complexity (number of parameters) and model adequacy (accommodating rate variation across lineages). Since the focus of our study is on the deep divergence times and the evolutionary rates, the rich fossil record in Early Cretaceous provided a great deal of information to produce reliable estimates deep in the tree. For the age estimates in the Late Cretaceous or later, biases might be severe due to limited fossil record. In particular, the divergence time of Anas and Gallus has been estimated at the early Eocene (Prum et al. 2015) associated with high evolutionary rates at the crown. In comparison, we inferred much younger age of 9.46 (4.57, 15.07) Ma with a long, low-rate ancestral branch. The underestimation was mainly due to lacking fossil or node calibrations within that clade, so the posterior estimate tended to be similar to the prior.

The gamma shape parameter (*α*) of character rate variation is larger than 1.0 (Table 1), indicating that the evolutionary rates are fairly homogeneous across characters. Note this gamma distribution models rate variation across characters and is independent of the gamma distributions for rate variation across branches in the relaxed clock model.

Partitioning the data is a common practice in molecular phylogenetic analyses (Nylander et al. 2004; Brown and Lemmon 2007). The different partitions may correspond to different genes and may also correspond to different codon positions in a protein-coding gene. On the other hand, morphological characters are typically treated as a single partition unless sufficient characters are available (Lee 2016) or simple model assumptions are made (Clarke and Middleton 2008). While successfully demonstrating the different evolutionary dynamics modeled by independent rate parameters in the six partitions, we note that the variance is very large (Fig. 2, widths of the error bars) due to limited number of characters in each partition. Further effort of coding more characters would refine the resolution.

In summary, the Bayesian tip dating approach implemented in MrBayes is a powerful and flexible tool to simultaneously estimate the tree topology, divergence times, evolutionary rates, and the other parameters of interest while accounting for their uncertainties. In the priors, we are able to incorporate the uncertainty of each fossil age, model the speciation, extinction, fossilization and sampling process explicitly, and take advantage of the relaxed clock model to investigate rate variation cross branches and partitions. It is feasible to integrate all available sources of information in the analysis, rather than discarding certain information or uncertainties in the parsimony and stepwise approach. Although the focal species are Mesozoic birds in this study, tip dating is a general framework applicable to a wide range of taxonomic groups with potential future extensions to the theoretical model and practical implementation.

### Availability

The models described above were implemented in MrBayes version 3.2.7 (Ronquist et al. 2012b. https://github.com/NBISweden/MrBayes; last accessed September 10, 2018).

## Supplementary Material

Data available from the Dryad Digital Repository: http://dx.doi.org/-.

## Acknowledgments

We sincerely thank three anonymous reviewers for constructive criticisms of the original article and excellent suggestions for improvement.

